# Pattern dynamics and stochasticity of the brain rhythms

**DOI:** 10.1101/2022.05.06.490960

**Authors:** C. Hoffman, J. Cheng, D. Ji, Y. Dabaghian

## Abstract

Our current understanding of brain rhythms is based on quantifying their instantaneous or time-averaged characteristics. What remains unexplored, is the actual structure of the waves—their shapes and patterns over finite timescales. To address this, we used two independent approaches to link wave forms to their physiological functions: the first is based on quantifying their consistency with the underlying mean behavior, and the second assesses “orderliness” of the waves’ features. The corresponding measures capture the wave’s characteristic and abnormal behaviors, such as atypical periodicity or excessive clustering, and demonstrate coupling between the patterns’ dynamics and the animal’s location, speed and acceleration. Specifically, we studied patterns of *θ* and *γ* waves, and Sharp Wave Ripples, and observed speed-modulated changes of the wave’s cadence, an antiphase relationship between orderliness and acceleration, as well as spatial selectiveness of patterns. Furthermore, we found an interdependence between orderliness and regularity: larger deviations from steady oscillatory behavior tend to accompany disarrayed temporal cluttering of peaks and troughs. Taken together, our results offer a complementary—mesoscale—perspective on brain wave structure, dynamics, and functionality.

## I. INTRODUCTION

Common approaches to studying “brain rhythms” can be broadly divided in two categories. The first is based on correlating the wave’s instantaneous phases, amplitudes and frequencies with parameters of cognitive, behavioral or neuronal processes. For example, instantaneous phases of *θ*-wave were found to modulate neuronal spikings [3, 4], while *γ*-waves’ amplitudes and frequencies link the synaptic [5–7] and circuit [8–10] current flow to learning dynamics. The second category of analyses is based on quantifying the brain waves’ time-averaged characteristics, e.g., establishing dependencies between the mean *θ*-frequency and the animal’s speed [11, 12] or acceleration [13] or linking rising mean *θ*- and *γ*-power to heightened attention states [13, 14] and so forth.

However, little work has been done to examine the waves’ overall shape, e.g., the temporal arrangement of peaks and troughs, or sequences of ripples or spindles generated over finite periods. Yet, the physiological relevance of the brain wave morphology is well recognized: rigidly periodic or excessively irregular rhythms that contravene a certain “natural” level of statistical variability are suggestive of circuit pathologies [15–21] or may indicate external driving [22, 23]. For example, the nearly periodic sequence of peaks shown on Fig. 1A is common for *θ*-waves, but certainly too orderly for the *γ*-waves. Conversely, the intermittent pattern on Fig. 1B is unlikely to appear among *θ*-waves, but may represent irregular *γ*-activity or typical sharp waves. On the other hand, the series of clumping peaks shown on Fig. 1C seem usual for in *γ* waves, but the cluttered pattern on Fig. 1D may be a manifestation of a particular process. In contrast, the waveforms shown on Fig. 1E,F appear to represent a mundane level of arhythmicity and temporal disorder expected in most *θ* and *γ* waves.

**FIG. 1.**
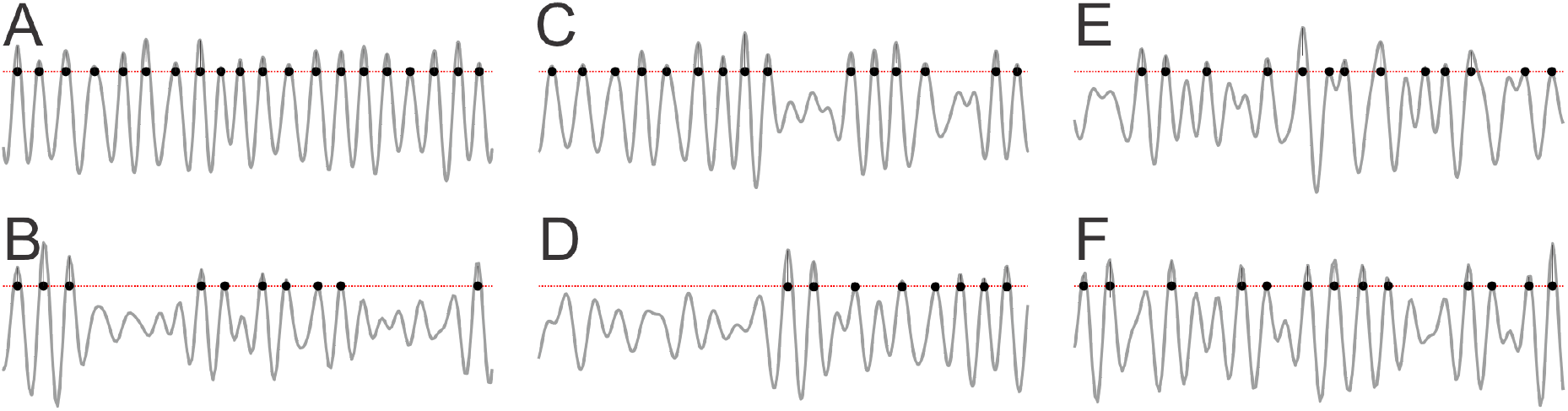
Waveform morphologies. **A**. A wave exhibiting a nearly periodic sequence of peak is commonly found among *θ*-oscillations, but rarely in higher-frequency waves. Conversely, the intermittent patterns shown on panel **B** may be exhibited by fast *γ* or sharp waves, but is atypical for lower-frequencies. The temporal clustering shown on panel **C** is not unusual for the *γ*-wave, but the uneven pattern on panel **D** would be an irregularity. In contrast, the patterns shown on panels **E** and **F** are all in all ordinary and seem to fluctuate with a fixed oscillatory rate. Shown are the peaks exceeding one standard deviation from the mean.

It remains unclear however, how to identify these patterns impartially, how to quantify the intuitive notions of “regularity,” “typicality,” “orderliness,” etc. Furthermore, since all brain waves exhibit a certain level of erraticness, it is unclear how justified are the experiential, visceral classifications of the waveforms. For example, might the “improbable” patterns illustrated above be attributed to mere fluctuations of otherwise regular waves, or should they be viewed as a structural peculiarity?

In the following, we address these questions in the context of two independent mathematical frameworks, using two cognate pattern quantifications that allow understanding the brain rhythms’ functional structure at intermediate timescales and their role in behavior and cognition.

## II. APPROACH

1. *Kolmogorov stochasticity, λ*, describes deviation of an ordered sequence, *X*, from the overall trend, and a remarkable observation made in [24] is that this score is universally distributed. As it turns out, deviations *λ*(*X*) that are too high or too low are rare: sequences with *λ*(*X*) ≤ 0.4 or *λ*(*X*) ≥ 1.8 appear with probability less than 0.3% (Fig. 2A,B, [25–30]). In other words, typical patterns are consistent with the underlying mean behavior and produce a limited range of *λ*-values, with mean *λ*^*^ ≈ 0.87. Thus, the *λ*-score can serve as a universal measure of *stochasticity*^1^ and be used for identifying statistical biases (or lack of thereof) in various patterns [31–36].
2. *Arnold stochasticity, β*, is alternative measure that quantifies whether the elements of a pattern “repel” or “attract” each other. Repelling elements seek to maximize separations and hence produce orderly, more equispaced arrangements, while attracting elements tend to cluster together. As shown in [37–40], for ordered patterns *β*(*X*) ≈ 1, for clustering ones *β*(*X*) can be high, while sequences with independent elements yield *β*-values close to the impartial mean, *β*^*^ ≈ 2 (Fig. 2C, Suppl. Sec. V). Thus, the *β*-score can be used to characterize *orderliness* of brain rhythms [37–40], complementing the *λ*-score.
3. *Time-dependence*. The recurrent nature of brain rhythms suggests dynamic generalization of *λ* and *β*. Given a time window, *L*, containing a sequence of events, *X*_*t*_, such as *θ*-peaks or Sharp Wave Ripples (SWR), evaluate the parameters *λ*(*X*_*t*_) and *β*(*X*_*t*_), then shift the window over a time step Δ*t*, evaluate the next *λ*(*X*_*t*+Δ*t*_) and *β*(*X*_*t*+Δ*t*_), and so on. The consecutive segments, obtained by small window shifts, *X*_*t*_, *X*_*t*+Δ*t*_, *X*_*t*+2Δ*t*_, …, differ only slightly from one another. The resulting time-dependencies *λ*(*t*) and *β*(*t*) will define the dynamics of stochasticity over the signal’s entire span.

**FIG. 2.**
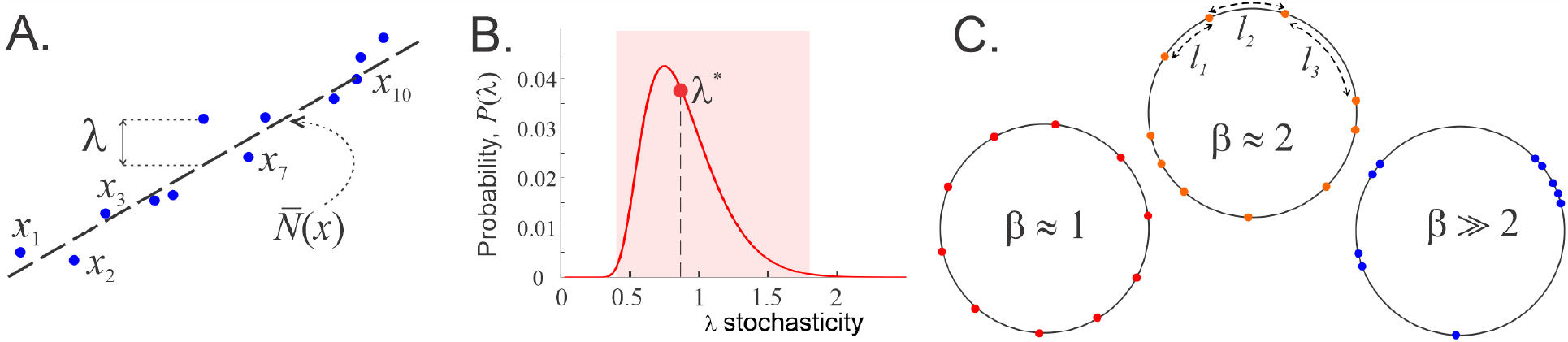
Stochasticity parameters. **A**. The elements of an ordered sequence *X* = {*x*_1_, *x*_2_, …, *x*_*n*_} following a linear trend 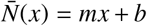(dashed line). The sequence’s deviations from the mean, *λ*(*X*), exhibit statistical universality and can hence impartially characterize the stochasticity of the individual data sequences *X*. **B**. The probability distribution of *λ*-scores is unimodal, with mean *λ*^*^ ≈ 0.87 (red dot). About 99.7% of sequences produce *λ*-scores in the interval 0.4 ≤ *λ*(*X*) ≤ 1.8 (pink stripe); these sequences are typical and consistent with the underlying mean behavior. In contrast, sequences with smaller or larger *λ*-scores are statistically uncommon. **C**. A sequence *X* arranged on a circle of length *L* produces a set of *n* arcs. The normalized quadratic sum of the arc lengths is small for orderly sequences, *β* ≈ 1 (left), as high as *β* ≈ *n* for the “clustering” sequences (right), and intermediate, *β* ≈ 2 (middle), for generic sequences.

For a visualization, one can imagine the elements of a given sample sequence, *X*_*t*+*k*Δ*t*_, as “beads” scattered over a necklace of length *L*. As the sliding window shifts forward in time, the beads shift back and may disappear at the back of the window, and new beads may appear toward the front, while a majority of the beads retain their relative positions. The corresponding *λ*- and *β*-values will then produce semi-continuous time-dependencies *λ*(*t*) and *β*(*t*) that quantify the “necklace dynamics”—gradual pattern changes. The parameter *β* then describes the orderliness of the beads’ distribution over the necklace, while *λ* measures how typical the beads’ arrangement is overall.

As an illustration, consider a data series *X* with random spacings between the adjacent values— intervals drawn from exponential, uniform, and Poisson distributions, with sample subsequences containing about 25 consecutive elements. As shown on Fig. 3A,C, the *λ*(*t*)-dependence of the exponential sequences remains, for the most part, constrained within the “typicality band” (pink stripe on Figs. 2B and Fig. 3A), while the uniformly distributed patterns are more variable and Poisson patterns follow the mean most closely. The *β*-scores of exponentially and uniformly distributed patterns are overall mundane, while the Poisson patterns exhibit periodic-like orderliness.

**FIG. 3.**
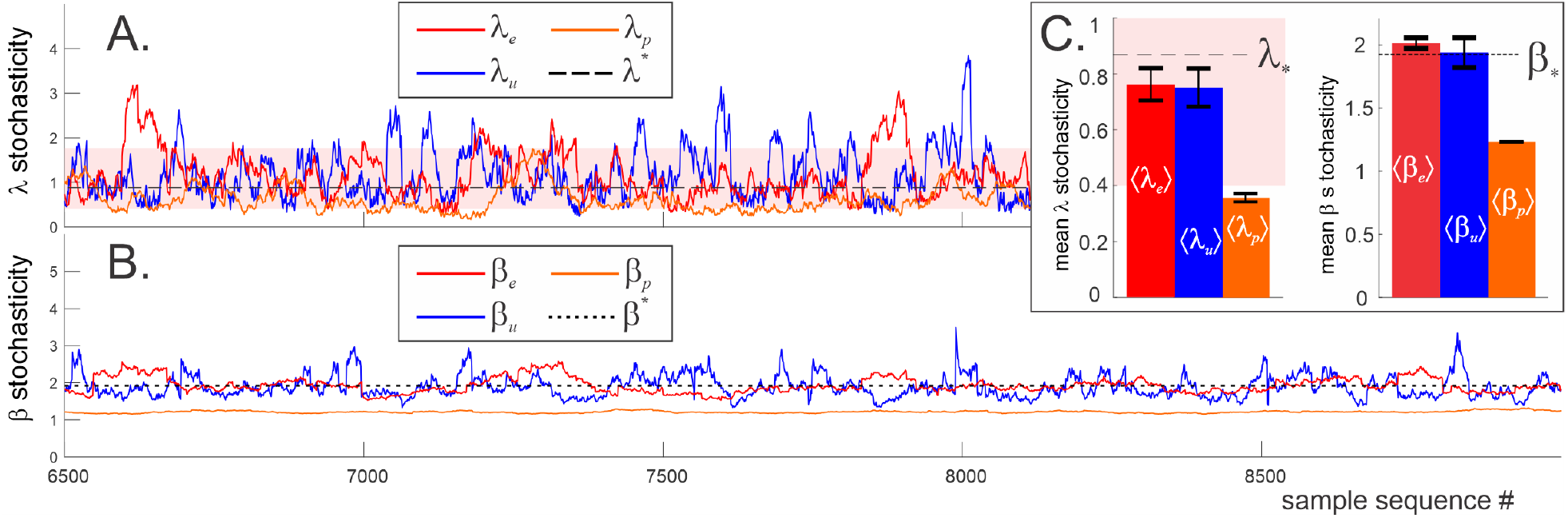
Pattern dynamics for three kinds of random sequences in which the intervals between consecutive points are distributed 1) exponentially with the rate *ν* = 2; 2) uniformly with constant density *ρ* = 1; or 3) with Poisson rate *μ* = 5. Sample intervals are selected proportionally to the distribution scales (*L*_*u*_ = 25*ρ, L*_*e*_ = 25*ν*, and *L*_*p*_ = 25*μ*, so that each sample sequence contains about *n* = 25 elements) and are shifted by a single data point at a time. **A**. The Kolmogorov parameter of the exponential sequence (red trace, *λ*_*e*_), uniform sequence (blue trace, *λ*_*u*_) and Poisson sequence (orange trace, *λ*_*p*_) remain mostly within the “pink zone” of stochastic typicality (same pink stripe as on Fig. 2B). *λ*_*u*_ is the most volatile and often escapes the expected range, whereas *λ*_*p*_ is more compliant, lingering below the expected mean *λ*_*p*_ ≲ *λ*^*^ ≈ 0.87 (black dashed line). **B**. The corresponding Arnold stochasticity parameters show similar behavior: *β*_*u*_ = 1.93 ± 0.2 fluctuates around the expected mean *β*^*^(25) = 1.92 (black dotted line). The exponential sequence has smaller *β*-variations and a slightly higher mean, *β*_*e*_ = 2 ± 0.04. The Poisson sequence is the least stochastic (nearly-periodic), with *β*_*p*_ = 1.22 ± 0.004, due to statistical suppression of small and large gaps. **C**. The mean stochasticity scores computed for about 10^4^ random patterns of each type.

The mean *λ*- and *β*-scores in the uniform and the exponential sequences are close to universal means, *λ*^*^ and *β*^*^, which shows that, on average, they are statistically unbiased. In contrast, the Poisson-distributed patterns are atypically orderly, due to statistically suppressed small and large gaps between neighboring elements (Fig. 3B).

The fluctuations of stochasticity scores—the rises and drops of *λ*(*t*) and *β*(*t*) dependencies on Fig. 3—are chancy, since random sequences vary sporadically between instantiations. In contrast, brain wave patterns may carry physiological information, and the dynamics of their stochasticity may serve as an independent characterization of circuit activity at a mesoscopic timescale in different behavioral and cognitive states, as discussed below.

## III. RESULTS

### A. Stochasticity in time

We analyzed Local Field Potentials (LFP) recorded in the hippocampal CA1 area of wild type male mice^2^ and studied their *θ*-wave, *γ*-wave, and SWR patterns [41]. The recurring nature of brain rhythms suggests that their key features distribute uniformly over sufficiently long periods. Therefore, the expected mean used for evaluating the Kolmogorov *λ*-parameter is linear,

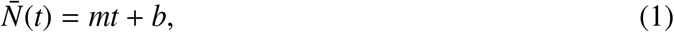

with the coefficients *m* and *b* obtained via linear regression. The lengths of the sample sequences were then selected to highlight the specific wave’s structure and functions, as described below.

**1. *θ*-waves** (4 − 12 Hz, [42, 43]) are known to correlate with the animal’s motion state, which suggests that the sample sequences should be selected at a behavioral scale [11–13]. In the analyzed experiments, the mice shuttled between two food wells on a U-shaped track, spending about 22 secs per lap (average for 5 mice, for both inbound and outbound runs) and consumed food reward over 17 secs (Fig. 4A). On the other hand, the intervals between successive *θ*-peaks distribute around the characteristic *θ*-period, 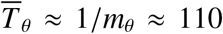 msecs, which defines the timescale of oscillatory dynamics (Fig. 4A). To accommodate both timescales, we used periods required to complete 1*/*6^th^ of the run between the food wells, *L*_*θ*_ ≈ 3.6 secs, containing about 20−30 peaks— large enough to produce stable *λ*- and *β*- scores [44–46], but short enough to capture the ongoing dynamics of *θ*-patterns.

**FIG. 4.**
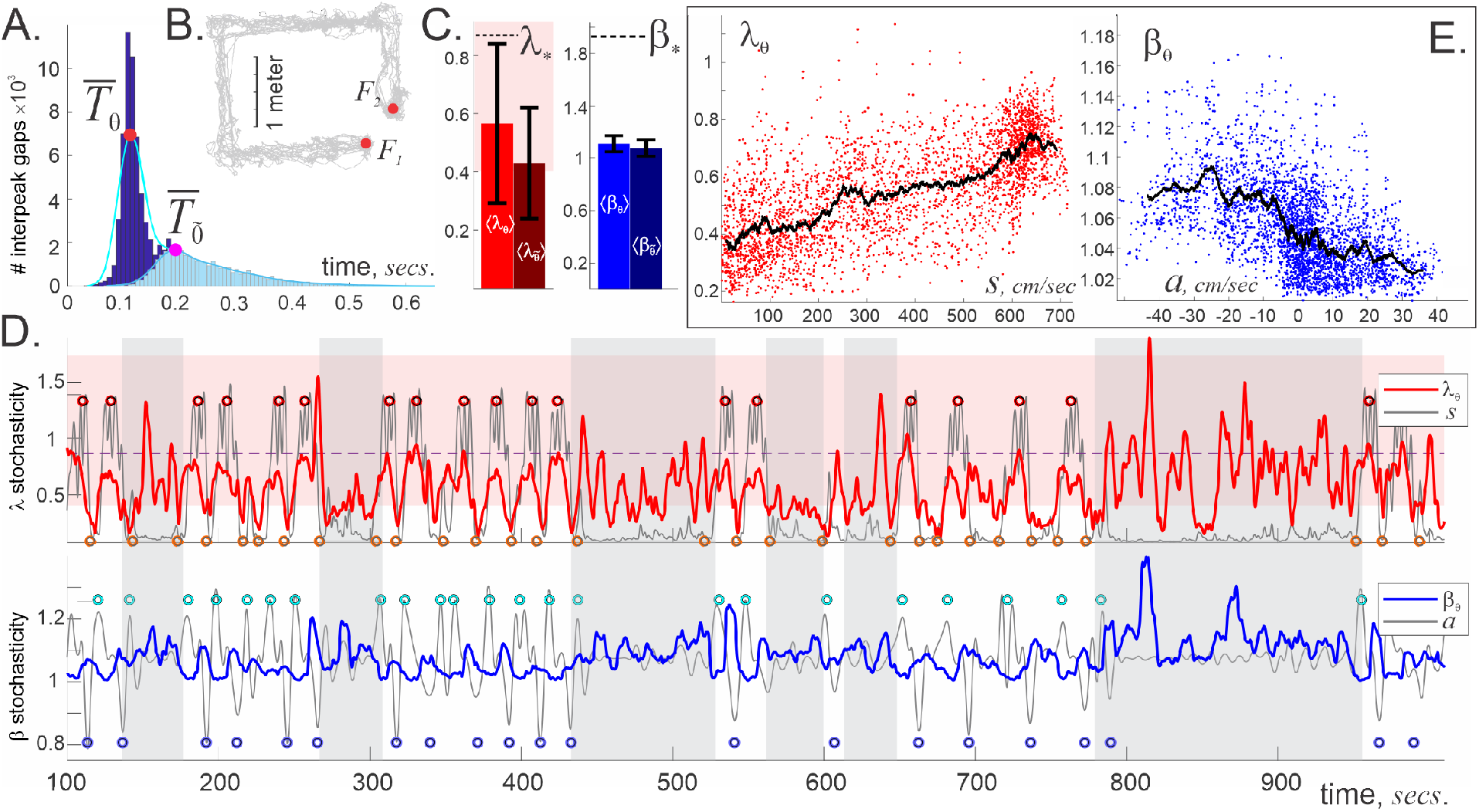
*θ*-wave’s stochasticity. **A**. A histogram of intervals between subsequent *θ*-peaks concentrates around the characteristic *θ*-period, 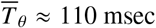 msec: gaps shorter than 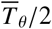 or wider than 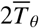 are rare. *θ*-amplitude, 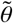, oscillates with 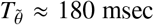 msec period. **B**. The animal’s lapses (trajectory shown by gray line) between food wells, *F*_1_ and *F*_2_, take on average 22 secs. **C**. Due to quasiperiodicity of the *θ*-wave and of its envelope, 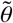, the average scores 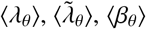 and 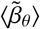 are significantly lower than the impartial means *λ*^*^ and *β*^*^, with small deviations (data for 5 mice). **D**. The dynamics of *λ*_*θ*_(*t*) (red trace, upper panel) correlates with the speed profile (gray line) when the mouse moves methodically. The *λ*_*θ*_(*t*)-stochasticity remains mostly within the “typical” range (pink stripe in the background), falling below it as the mouse slows down. For rapid moves there is a clear similarity between the *λ*_*θ*_-score and the speed, e.g., their peaks and troughs roughly match. When the mouse meanders (vertical gray stripes), the coupling between speed and *λ*_*θ*_-stochasticity is lost. The Arnold score *β*_*θ*_(*t*) (blue trace, lower panel) remains close to *β*_min_ = 1, affirming *θ*-wave’s quasiperiodicity. Note the antiphasic relationship between the *β*_*θ*_-stochasticity and the acceleration *a*(*t*) (the latter graph is shifted upwards to match the mean level ⟨*β*_*θ*_⟩): *θ*-periodicity loosens as the animal slows down (*β*_*θ*_- splashes correlate with animal’s deceleration) and sharpens as he speeds up. **E**. Locally averaged 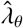-score grows with speed, whereas 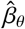 tends to drop down with acceleration.

The resulting mean Kolmogorov score ⟨*λ*_*θ*_⟩ = 0.54 ± 0.12 is low, indicating that, on average, *θ*-cycles closely follow the prescribed trend (1). The mean Arnold score ⟨*β*_*θ*_⟩ = 1.1 ± 0.03 ≈ *β*^min^ also points at the near-periodic behavior of the *θ*-wave (Fig. 4C). Nonetheless, *θ*-patterns exhibit complex dynamics that differ between quiescence and active states and couple to the animal’s speed and acceleration.

#### Fast moves

As mentioned above, the experimental design enforces recurrent behavior, in which speed goes up and down repeatedly as the animal moves between the food wells (Fig. 4B). When the mouse moves methodically (lap time less than 25 sec), *λ*_*θ*_ rises and falls along with the speed with surprising persistence (Fig. 4D). Yet, the *θ*-patterns appearing in this process are stochastically generic—the entire sequence of *λ*_*θ*_-values remains mostly within the “domain of stochastic typicality” (pink stripe on Figs. 4C, D and 2B), below the universal mean *λ*^*^. However, the patterns become overtly structured as the animal slows down, when the Kolmogorov scores drop below *λ*_*θ*_ ≈ 0.1, exhibiting uncommon compliance with the mean behavior. Such values of *λ*_*θ*_ can occur by chance with vanishingly small probability Φ(0.1) *<* 10^−17^ (see formula (5) in Sec. V), which, together with small *β*_*θ*_-score, *β*_*θ*_ ≈ 1, imply that limited motor driving reduces *θ*-wave to a simple nearly-harmonic oscillation with a base frequency *ν* ≈ 8 Hz.

Increasing speed randomizes *θ*-patterns: the faster the mouse moves, the higher the *λ*_*θ*_-score. Furthermore, the shape of *λ*_*θ*_(*t*) dependence exhibits an uncanny resemblance to the speed profile *s*(*t*) (Fig. 4D). To quantify this effect, we used the Dynamic Time Warping (DTW) technique that uses a series of local stretches to match two functions—in this case, *λ*_*θ*_(*t*) and *s*(*t*)—so that the net stretch can be interpreted as separation, or distance^3^ between functions in “feature space” [47, 48]. In our case, the DTW-distance between the speed *s*(*t*) and Kolmogorov score *λ*_*θ*_(*t*) during active moves is small, *D*(*λ*_*θ*_, *s*) = 19.6%, indicating that the deviations of the *θ*-patterns from the mean reflect the animals’ mobility (SFig. 1).

Note that DTW-affinity between *λ*_*θ*_(*t*) and speed does not necessarily imply a direct functional dependence between these quantities. Indeed, plotting points with coordinates (*s, λ*_*θ*_) yields scattered clouds, suggesting a broad trend, rather than a strict relationship (Fig. 4E). However, if the *λ*-scores and the speeds are *locally averaged*, i.e., if each individual *s*- and *λ*-value is replaced by the mean of itself and its adjacents, then the pairs of such *local means*, 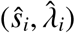), reveal a core dependence: increasing speed of the animal entails higher variability of the *θ*-patterns.

In the meantime, the Arnold stochasticity score, *β*_*θ*_(*t*), is closely correlated with the mouse’s acceleration, *a*(*t*). As shown on Fig. 4D, the *β*_*θ*_-score rises as the mouse decelerates (*θ*-wave clumps) and falls when he accelerates (*θ*-wave becomes more orderly), producing a curious antiphasic *β*_*θ*_-*a* relationship, which is also captured by the local averages 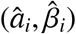 (Fig. 4E). The distance between *β*_*θ*_(*t*) and −*a*(*t*) (the minus sign accounts for the antiphase) is *D*(*β*_*θ*_, −*a*) = 33.6%, which means that speed influences the *θ*-wave’s statistical typicality more than acceleration impacts its orderliness.

#### Slow moves

When the mouse meanders and slows down (lapse time over 25 sec), *θ*-patterns change: the *λ*_*θ*_-score increases in magnitude and uncouples from speed, (DTW distance is twice that of the fast moves case, *D*(*λ*_*θ*_, *s*) = 38.8%), suggesting that, without active motor driving, *θ*-rhythmicity is less controlled by the mean oscillatory rate, i.e., is more randomized. The Arnold’s parameter *β*_*θ*_ also slightly increases, *D*(*β*_*θ*_, −*a*) = 36.1%, indicating concomitant *θ*-disorder.

Overall, the combined *λ*_*θ*_-*β*_*θ*_ dynamics suggests that during active behavior, the shape of the *θ*-wave is strongly controlled by the mouse’s moves. Highly ordered, nearly periodic *θ*-peaks appear when the animal starts running—the *θ*-frequency range then narrows to the mean, expected value. The increasing speed stirs up the *θ*-patterns; the disorder grows and reaches its maximum when the animal moves fastest and begins to slow down. During periods of inactivity, the coupling between *θ*-patterns and speed is weakened and then reinforced as the mouse stiffens his resolve.

We emphasize however, that these dependencies should not be viewed as naïve manifestations of known couplings between instantaneous or time-averaged frequency with the animals’ speed or acceleration [12, 13]. Indeed, the *θ*-frequency alters at the same rate as the *θ*-amplitude—many times over the span of each *θ*-pattern, providing an instantaneous characterization of the wave [50, 51]. In contrast, the stochasticity parameters describe the wave form as a single entity (Fig. 4A,C). On the other hand, time-averaging levels out fluctuations and highlights mean trends, whereas Kolmogorov and Arnold parameters are sensitive to individual elements in the data sequences. *Thus, λ and β scores describe of wave shapes without defeaturing, putting each pattern, as a whole, into a statistical perspective*. It hence becomes possible to approach questions addressed in the Introduction: identify typical and atypical wave patterns, quantify levels of their orderliness, detect deviations from natural behavior and so forth.

**2. *γ*-waves** (30 −80 Hz, [7]) exhibit a wider variety of patterns than *θ*-waves. The interpeak intervals between consecutive *γ*-peaks, *T*_*γ*_, are nearly-exponentially distributed, which implies that both smaller and wider *γ*-intervals are statistically more common (Fig. 5A).

**FIG. 5.**
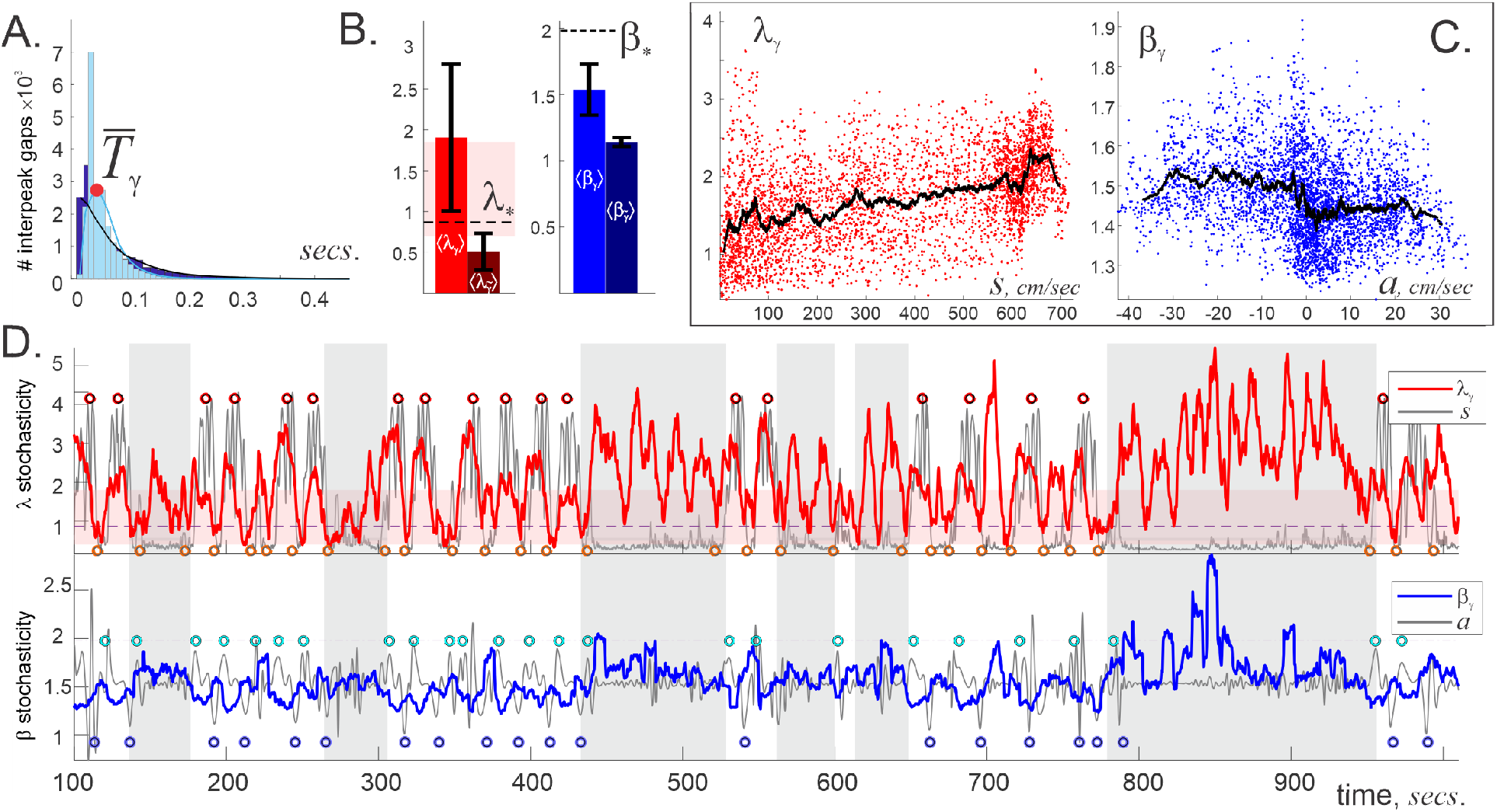
*γ*-wave stochasticity. **A**. A histogram of *γ*-interpeak intervals exhibits an exponential-like distribution with mean characteristic *γ*-period, 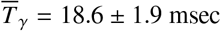 msec, about six times smaller than 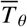. **B**. The average scores ⟨*β*_*γ*_⟩ and ⟨*λ*_*γ*_⟩ are higher than for the *θ*-wave, indicating that *γ*-patterns are more diverse than *θ*-patterns. **C**. Locally averaged 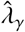-score grows with speed, while 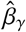 switches from higher to lower level with increasing acceleration. **D**. The dynamics of the *λ*_*γ*_-score (top panel) correlates with changes in the speed when the animal moves actively. Note that *λ*_*γ*_ often exceeds the upper bound of the “pink stripe,” i.e., *γ*-waves often produce statistically uncommon patterns, especially during rapid moves. The *β*_*γ*_-score (bottom panel) correlates with the animal’s acceleration, which is lost when lap times increase (gray stripes).

#### Fast moves

For consistency, the sample sequences, *X*_*γ*_, were drawn from the same time windows, *L*_*γ*_ = *L*_*θ*_ ≈ 6 secs, which contained, on average, about 300 elements that yield a mean Kolmogorov score ⟨ *λ*_*γ*_⟩ = 1.84 ± 1.03—more than twice higher than the impartial mean *λ*^*^ and three times above the ⟨ *λ*_*θ*_⟩ score. Such values can randomly occur with probability 1 − Φ(1.84) ≲ 2 · 10^−3^, which suggests that generic *γ*-patterns are statistically atypical and may hence reflect organized network dynamics, rather than random extracellular field fluctuations (Fig. 5B). The average Arnold parameter also grows compared to the *θ*-case, but remains lower than the impartial mean, ⟨*β*_*γ*_⟩ = 1.61 ± 0.53 *< β*^*^, implying that, although *γ*-waves are more disordered than the *θ*-waves, they remain overall oscillatory.

On average (for all five mice), the Kolmogorov score, *λ*_*γ*_(*t*), escapes the domain of “stochastic typicality” approximately half of time through transitions that closely follow speed dynamics. Together with the observation that *λ*_*γ*_ takes a larger range of values than *λ*_*θ*_ (*γ*-patterns deviate more from the average as speed increases), this suggests that there is a greater diversity of *γ*-responses to movements (Fig. 5C). In particular, the dependence between locally averaged 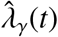 and *ŝ*(*t*) is less tight: the average DTW distance, *D*(*λ*_*γ*_, *s*) ≈ 23.4%, is slightly higher than the distance in the *λ*_*θ*_(*t*) dynamics, which illustrates that *γ*-patterns are less sensitive to speed than *θ*-patterns.

From a structural, *β*-perspective, *γ*-wave becomes closer to periodic when an actively moving animal slows down: during these periods, *β*_*γ*_-score reduces close to its minimal value, when the corresponding Kolmogorov score also drops to *λ*_*γ*_ ≈ 0.2. Since the latter is unlikely to occur by chance (cumulative probability of that is Φ(0.2) ≲ 10^−12^), these changes may represent structured network dynamics. The highest deviations of *γ*-patterns from the mean (*λ*_*γ*_ 3) are accompanied by high *β*_*γ*_-scores, which happen as the mice slow down from maximal speed and implies that circuit activity is least structured during these periods (Fig. 5D). The relation between *γ*-orderliness, *β*_*γ*_(*t*), and acceleration is also similar to the corresponding dependence in the *θ*-case: acceleration induces stricter *γ*-rhythmicity and deceleration randomizes *γ*-patterns, with about the same overall DTW distance, *D*(*β*_*γ*_, −*a*) ≈ 34.4%.

*During slower movements*, the *γ*-dynamics change qualitatively: the magnitudes of both *λ*_*γ*_(*t*) and *β*_*γ*_(*t*) grow higher, indicating that decoupling from motor activity enforces statistically atypical *γ*-rhythmicity in the hippocampal network, as in the *θ*-waves. In particular, the uncommonly high *β*_*γ*_ scores point at frequent *γ*-bursting during quiescence.

Once again, we emphasize that these results do not represent known correlations between instantaneous or time-averaged *γ*-characteristics and motion parameters [52, 53]. Rather, the outlined *λ*_*γ*_(*t*) and *β*_*γ*_(*t*) dependencies capture pattern-level dynamics of *γ*-waves that reflect circuit activity at an intermediate timescale. As an illustration, note that the amplitude of *γ*-waves, 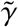, along with the instantaneous *γ*-frequency, *ω*_*γ*_, have low stochasticity scores, comparable to the ones produced by the Poisson process, 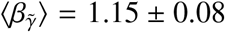 and 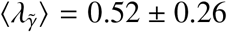 (Fig. 3). Thus, although instantaneous characteristics exhibit restrained, quasiperiodic behavior, they allow a rich morphological variety of the underlying *γ*-oscillations.

**3. Sharp Wave Ripples** (SWRs), the high amplitude splashes (over 2 − 3 standard deviations from the mean) of high frequency waves (150−250 Hz, [41]), exhibit the richest pattern dynamics.

*During fast moves*, SWR events appear at approximately the same exponential rate as the 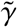-peaks, 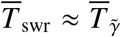 msec, but exhibit higher *λ*-scores, ⟨*λ*_swr_⟩ = 2.40 ± 1.57, over the same sampling periods *L*_swr_ ≈ 6 sec (Fig. 6A,B). The low probability of these patterns (1 − Φ(2.5) ≲ 10^−6^) and the relatively high mean *β*-score, ⟨*β*_swr_⟩ = 1.71 ± 0.64, indicate that SWRs tend to exhibit intermittent clustering that may reflect brisk, time-localized circuit activity, such as rapid replays and preplays of the hippocampal place cells [54–60].

**FIG. 6.**
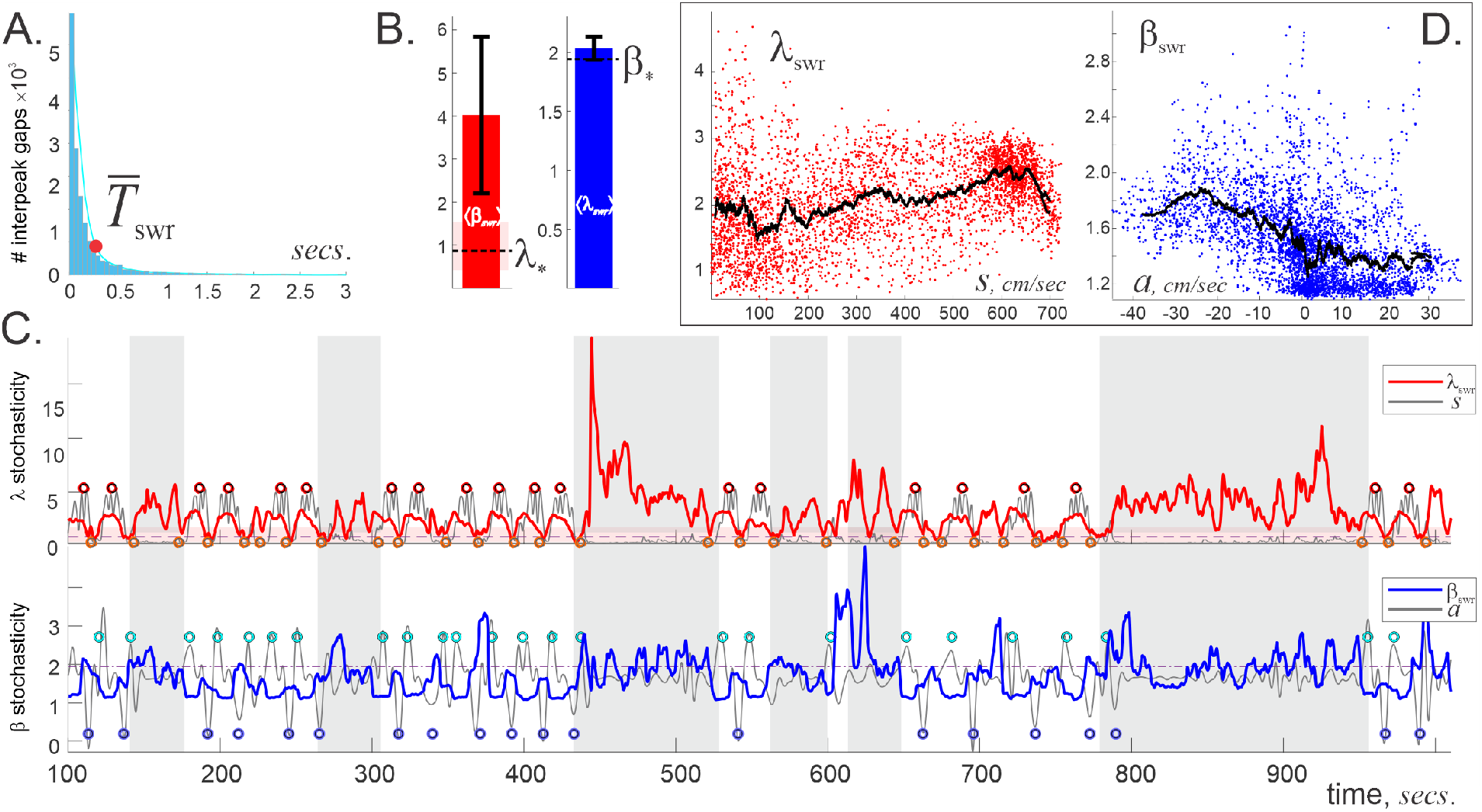
Sharp Wave-Ripples’ stochasticity. **A**. A histogram of intervals between SWR events is nearly exponential. **B**. The averages ⟨*λ*_swr_⟩ and ⟨*β*_swr_⟩ are high, indicating both frequent deviation of SWR events from the mean and higher temporal clustering than for the *θ* and *γ*-patterns. **C**. The animal’s speed (gray line, top panel) correlates with the Kolmogorov parameter *λ*_swr_ during fast exploratory lapses. During inactivity (vertical gray stripes) the *λ*_swr_-stochasticity uncouples from speed, exhibiting high spikes that mark strong “fibrillation” of SWR patterns. The antiphasic relationship between the animal’s acceleration *a*(*t*) (gray line, bottom panel) and Arnold score *β*_swr_(*t*) shows that SWRs tend to cluster when as the animal decelerates, while acceleration enforces periodicity. During slower moves (gray stripes), the relationship between speed, acceleration, and stochasticity is washed out and stochastically improbable patterns dominate. **D**. Locally averaged 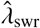 grows with speed and 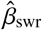 drops with acceleration.

Interestingly, SWR-patterns also correlate with the animal’s speed profile about as much as *γ*-patterns, *D*(*λ*_swr_, *s*) ≈ 23.8% [58]. The *β*_swr_(*t*)-dependence displays the familiar antiphasic relationship with the animal’s acceleration—SWR events tend to cluster more when the animal slows down (Fig. 6C,D). However, orderliness of SWRs is driven by acceleration stronger than orderliness of *γ*-patterns: the range of *β*_swr_-scores is twice as wide as the range of *β*_*γ*_-scores (broader pattern variety), with a similar DTW distance *D*(*β*_swr_, −*a*) ≈ 39.7%.

*During quiescent periods*, both *λ*_swr_ and *β*_swr_ grow and exhibit extremely high spikes, indicating that endogenous network dynamics produce stochastically improbable, highly clustered SWR sequences. Physiologically, these statistically uncommon SWR patterns may indicate sleep or still wakefulness replay activity, known to play an important role in memory consolidation [61, 62].

Overall, the temporal clumping comes forth as a characteristic feature of the SWR events, suggesting that SWRs are manifestations of fast, targeted network dynamics that brusquely “ripple” the extracellular field, unlike the rhythmic *θ* and *γ*-undulations [41].

### B. Stochasticity in space

Distributing the *λ* and *β* scores along the animal’s trajectory yields *spatial maps of stochasticity* for each brain rhythm and reveals a curious spatial organization of LFP patterns with similar morphology. As shown on Fig. 7, higher *λ*-values for all waves are attracted to segments where the mouse is actively running with maximal speed, furthest away from the food wells. Patterns that are close to the expected average (low-*λ*) concentrate in the vicinity of food wells where the animal moves slowly. The latter domains also tend to host high *β*-scores that appear as the animal approaches the food wells, as well as the lowest *β*s, which appear as the animal accelerates away [59]. In other words, the LFP waves become more “trendy” and, at the same time, more structured (either more periodic or more clustered) over the behaviorally important places (e.g., food wells) that require higher cognitive activity. On the other hand, the outer parts of the track, where the brain waves are less controlled by the mean and remain moderately disordered, are marked by uncommon patterns.

**FIG. 7.**
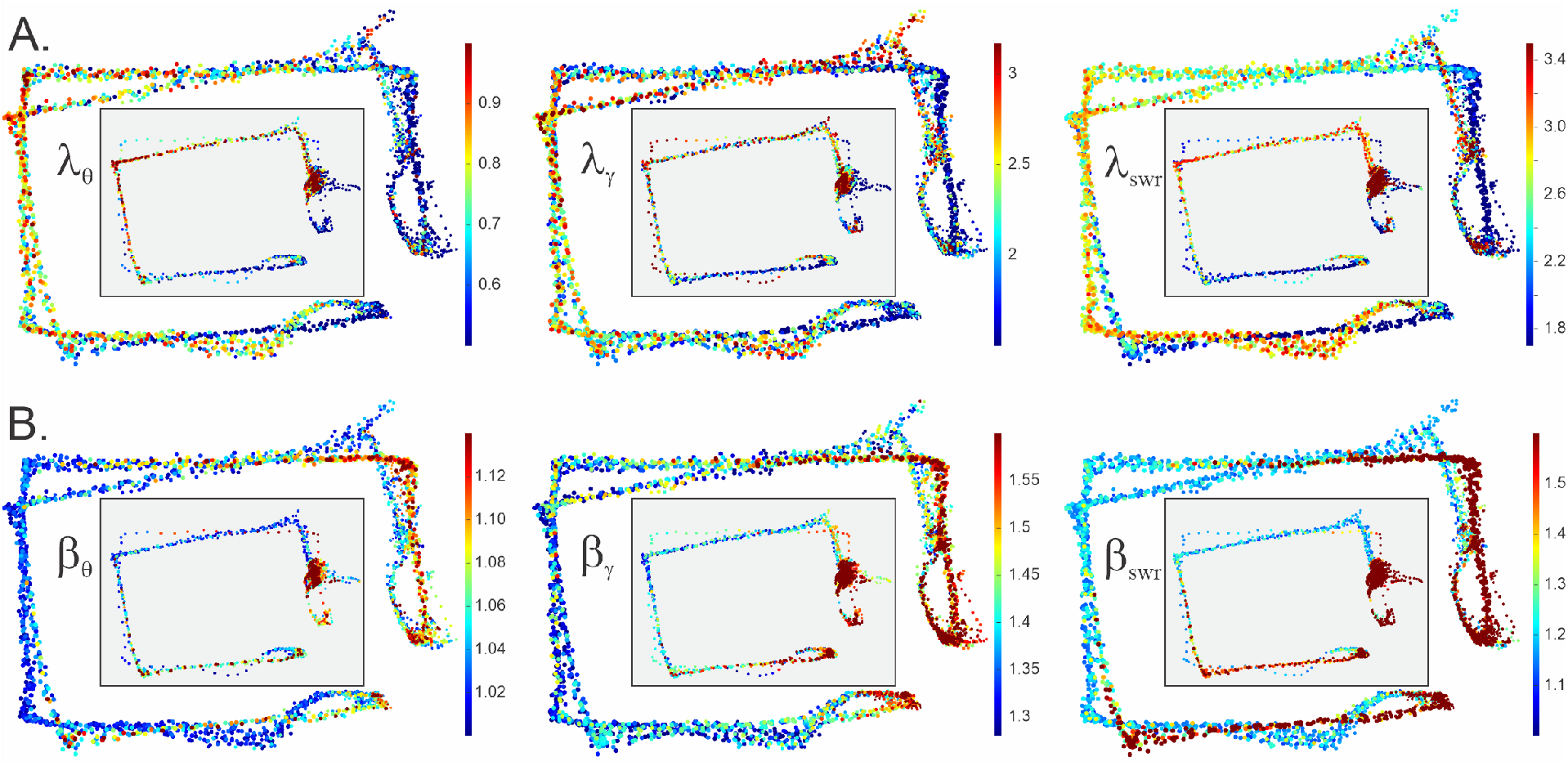
Spatial stochasticity maps were obtained by plotting *λ* and *β* parameters along the trajectory. **A**. The *λ*-maps show that *θ*-wave, *γ*-wave and SWRs generally follow the mean trend near the food wells (with scattered wisps of high stochasticity) and deviate from the mean mostly over the areas most distant from the food wells. The smaller maps in the gray boxes represent slow lapses: the overall layout of high-*λ* and low-*λ* fields is same as during the fast moves, which suggests spatiality of *λ*-stochasticity. **B**. The behavior of *β*_*θ*_ is opposite: the “uneventful,” distant run segments attract nearly-periodic behavior, while the food wells attract time-clumping wave patterns.

Intriguingly, the same map structure is reproduced during slow lapses, when the motor control of the patterns weakens, suggesting that speed and acceleration are not the only determinants of the LFP patterns. As shown on Fig. 7, even when the mouse dawdles, the waves tend to deviate from the mean around the outer corners and follow the mean in the vicinity of the food wells. Similarly, the patterns start clumping as the mouse approaches the food wells, and distribute more evenly as he moves away.

These results suggest that spatial context may, by itself, influence hippocampal brain rhythm structure, which is reminiscent of the place-specific activity exhibited by spatially tuned neurons, e.g., place cells [63] or parietal neurons [64]. For example, the “bursting” (high-*β*) fields and “domains of evenness” (small *λ*) surround food wells; the quasiperiodicity fields (small *β*s) as well as “wobbling-waves” (large *λ*s) stretch over the outer segments (Fig. 7). Physiologically, this “spatiality of stochasticity” may reflect a coupling between the hippocampal place-specific spiking activity and extracellular field oscillations.

### C. *λ*-*β* relationships

The definitions of the stochasticity scores *β* and *λ* do not imply an *a priori β*-*λ* relationship. Indeed, plotting sets of points with coordinates (*β, λ*) for all sample sequences, *X*_*θ*_, *X*_*γ*_ and *X*_swr_, yields scattered clouds, rather than curve-like graphs (Fig. 8A). However, computational studies of number-theoretic sequences carried in [37–40] suggest that such behavior may be caused by the occasional large contributions from atypical sequences and that local smoothing may yield much tighter couplings between the stochasticity parameters.

**FIG. 8.**
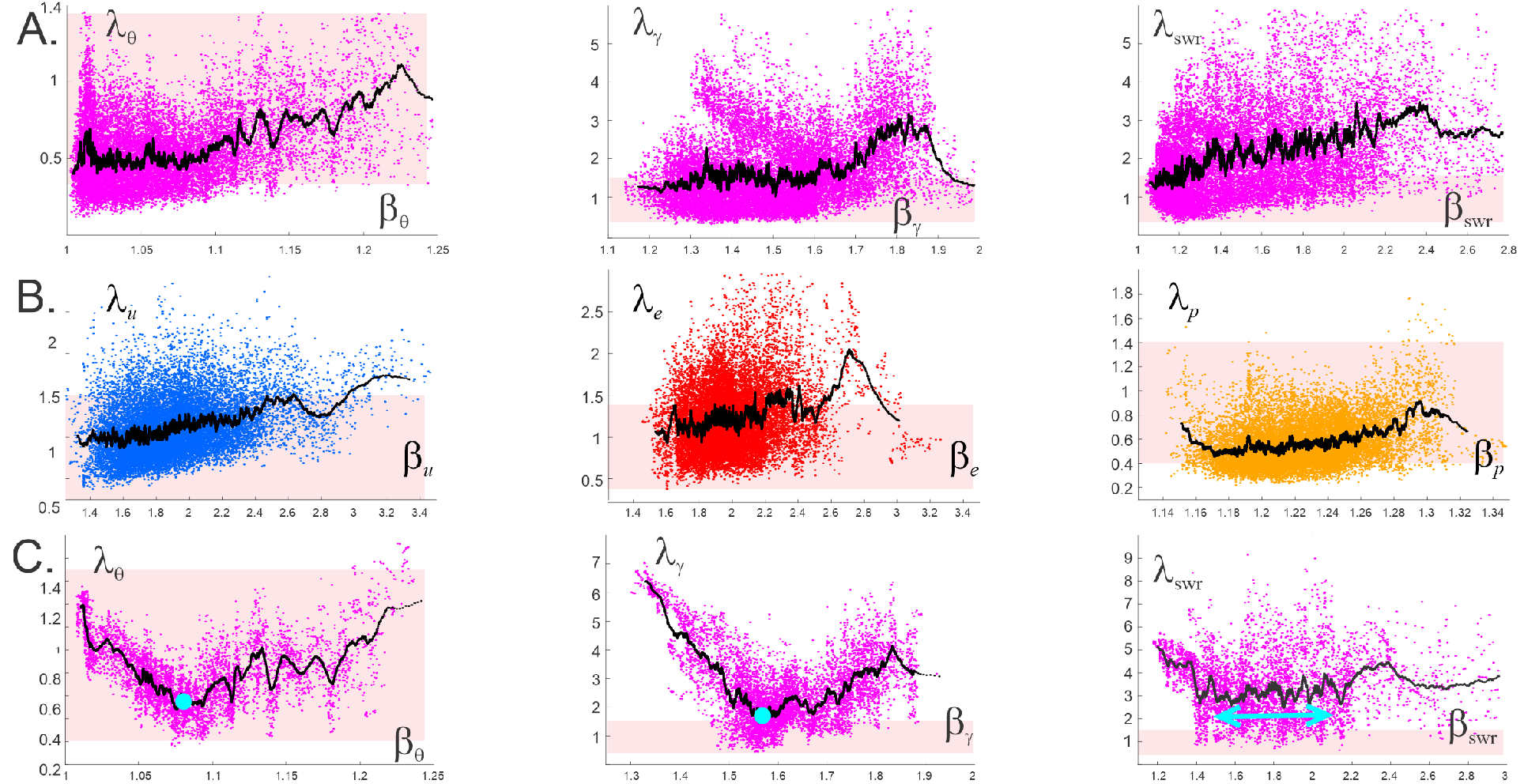
Dependencies between stochasticity parameters. **A**. Points with coordinates (*β*_*i*_, *λ*_*i*_) computed for each individual sample sequence produce clouds that imply no strict *β*(*λ*) dependence. However, locally averaged stochasticity parameters 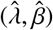 exhibit much tighter relationships (black dots). The growing 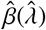 trends indicate that ordered, semiperiodic sequences (low 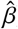) tend to accompany samples that comply with the expected behavior (low 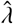). Conversely, patterns tend to fibrillate more as they deviate farther from the mean and abandon the pink stripe of “stochastic typicality.” **B**. The (*β, λ*) pairs for sequences drawn from the three random distributions (Fig. 3) produce similar clouds (uniform, blue dots; exponential, red dots; Poisson, orange dots). Locally averaged scores indicate tight 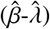 relations for all these series trend similarly to the brain waves’ local averages. **C**. The 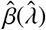 trends evaluated for individual mice may exhibit non-generic features. In this case, the oscillatory rate of semi-periodic *θ*-, *γ*- and SWR sequences (low 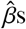) deviates significantly from the predicted mean (large 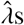). As the disorder increases, the oscillatory rate gets closer to the predicted mean, reaching the minimal 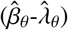 and 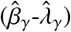 combinations (cyan dot) before retaking a joint growth trend. In case of the SWRs, the minimum is spread into a plateau where changes in 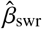 do not affect 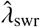.

To study whether similar phenomena take place in the LFP sequences, we evaluated the the local averages 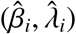, which revealed 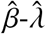 relationships illustrated on Fig. 8. For the full dataset that includes all mice, we observe that orderly sequences (low 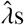-scores) tend to follow the prescribed mean trend more closely (low 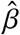). In contrast, patterns that deviate from the expected mean tend to produce disordered and clumping sequences, notably for *γ*-waves and SWRs.

Curiously, similar 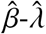 behavior are exhibited by random sequences (uniformly, exponentially and Poisson-distributed, Fig. 8B), which suggests that the tendency of *λ* to rise with growing *β*, observed in large volumes of heterogeneous data (different mice) may not be of a specifically physiological nature, but may reflect mathematical connections between the stochasticity indices [28–30].

In view of these results, it is surprising that individual mice can exhibit personalized dependencies between predictability, *λ*, and orderliness, *β*, of their LFP patterns. In the case illustrated on Fig. 8, nearly periodic (small 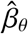) *θ*-patterns tend to deviate from the predicted behavior quite significantly. As the disorder increases, the patterns become more compliant with the underlying mean, until a trough of 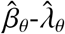 dependence is reached. Then the tendency is reversed: growing *θ*-disorder is accompanied by further deviation from the mean. Note that the entire stochasticity dynamics remain within the “typicality zone,” 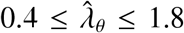. Analogous behavior is exhibited by the *γ*-wave but at a larger scale: orderly *γ*-waves, 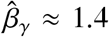, are concomitant with *λ*-scores as high as *λ*_*γ*_ ≈ 6.5 (highly improbable patterns, 1 − Φ(6.5) ; ≲ 10^−50^). As 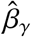 grows, 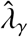 decreases, approaching stochastic commonality at the lowest point. Then, as 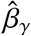 grows further, the disordered sequences increasingly deviate from the expected mean. The 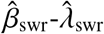 dependence is less tight but exhibits a similar trend: at first, small 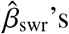 pair with higher 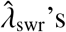, then level out over an intermediate range of *β*’s (the minimum is flattened out in contrast with *θ*- and *γ*-waves), and then grows again with the increasing SWR-disorder.

Importantly, we found no relationship between the *λ*_*θ*_, *λ*_*γ*_, and *λ*_*swr*_, nor between *β*_*θ*_, *β*_*γ*_, and *β*_*swr*_ (SFig. 2), which implies that the *λ*_*_ and the *β*_*_-scores associated with different waves are largely independent, providing their own, autonomous characterizations of the wave shapes.

## IV. DISCUSSION

The recorded LFP signals are superpositions of locally induced extracellular fields and inputs transmitted from anatomically remote networks. The undulatory appearance of the LFP is often interpreted as a sign of structural and functional regularity^4^, but the dynamics of these oscillations is actually highly complex. Understanding the balance between deterministic and stochastic components in LFPs, as well as questions about their continuity and discreteness pose significant conceptual challenges, as it happened previously in other disciplines^5^.

Structurally, LFP rhythms may be described through discrete sequences of wave features (heights of peaks, specific phases, interpeak intervals, etc.), or viewed as transient series—bursts— of events, as in the case of SWRs [65]. It is well recognized that such sequences are hard to decipher and to forecast, e.g., a recent discussion of a possible role of bursts in brain waves’ genesis posits: *“An important feature that sets the burst scenario apart is the lack of continuous phase-progression between successive time points—and therefore the ability to predict the future phase of the signal—at least beyond the borders of individual bursts”* [65]. In other words, the nonlinearity of LFP dynamics, as well as its transience and sporadic external driving, result in effective stochasticity of LFP patterns—an observation that opens a new round of inquiries [65, 66]. For example, how exactly should one interpret the “unpredictability” of a temporal sequence? Does it mean that its pattern cannot be resolved by a particular algorithm, or that it is unpredictable in principle, “genuinely random,” such as a gambling sequence? What is the difference between the two? How is the apparent randomness of LFP rhythms manifested physiologically? Are the actual network computations based on “overcoming the apparent randomness” and somehow deriving the upcoming phases or amplitudes from the preceding ones, or may there be alternative ways of extracting information? Does the result depend on the “degree of randomness” and if so, then how to distinguish between a “more random” and a “less random” patterns? These questions are not technical, pertaining to a specific mechanism, nor specifically neurophysiological; rather, these are fundamental problems that transcend the field of neuroscience. Historically, similar questions have motivated mathematical definitions of randomness that are still debated to this day [69–71].

One approach to addressing these issues was suggested by Kolmogorov in 1933 (also the year when brain waves were discovered [72]), based on the statistical universality of stochastic deviations from the expected behavior [24, 25]. From Kolmogorov’s perspective, randomness is contextual: if a sequence *X* deviates from an expected mean behavior within bounds established by the distribution (3), then *X* is *effectively* random. In other words, an individual sequence may be viewed as random if it could be randomly drawn from a large pool of similarly trending sequences, with sufficiently high probability. This view permits an important conceptual relativism: even if a sequence is produced by a specific mechanism or algorithm, it can still be viewed as random as long as its *λ*-score is “typical” according to the statistics (3). For example, it can be argued that geometric sequences are typically more random than arithmetic ones, although both are defined by explicit formulae [26–30]. By analogy, individual sample sequences of *θ, γ*, or SWR events may be generated by specific synchronization mechanisms at a precise timescale, and yet they may be empirically classified and quantified as stochastic.

A practical advantage of Kolmogorov’s approach is that mean trends, such as (1), can often be reliably established, interpreted, and then used for putting the stochasticity of the underlying sample patterns into a statistical perspective. Correspondingly, assessments based on *λ*-scores were previously applied in a variety of disciplines from genetics [31–33] to astronomy [34], and from economics [35] to number theory [26–30, 36–40, 73]. Some work has also been done in brain wave analyses, e.g., for testing normality of electroencephalograms’ long-term statistics [74–77]. Arnold’s *β*-score provides an independent assessment of orderliness (whether elements of an arrangement tend to attract, repel or be independent of each other) and it has not been, to our knowledge, previously used in applications.

Shifting window analyses ground the *λ*- and *β*-values in the context of preceding and upcoming observations. Since neighboring patterns change only marginally, the time-dependent *λ*(*t*) and *β*(*t*) describe quasi-continuous pattern dynamics at a *mesoscale*—over the span of several undulations—which complements the microscale (instantaneous) and macroscale (time-averaged) assessments. While the individual, “stroboscopically selected” patterns can be viewed as stochastic, the continuous *λ*(*t*) and *β*(*t*) dependencies describe ongoing pattern dynamics.

Importantly, Kolmogorov’s and Arnold’s scores are impartial and independent from physiological specifics or contexts, thus providing self-contained semantics for describing the LFP data and a novel venue for analyzing the underlying neuronal mechanisms. It becomes possible to distinguish “statistically mundane” LFP patterns from exceptional ones and to capture the transitions between them, as well as to link pattern dynamics to changes in the underlying network’s dynamics (Figs. 4-7). For example, since *θ*-bursts are physiologically linked to long-term synaptic potentiation [78, 79], *θ*-patterns with high *β*-scores may serve as markers of plasticity processes taking place in the hippocampal network at specific times and places [80–82]. Furthermore, high-*β*_*θ*_ regions near food wells indicate that reward proximity may trigger hippocampal plasticity and, since hippocampal neurons’ spiking is coupled to *θ*-cycles [3], have a particular effect on memory (Fig. 7). On the other hand, low-*β*_*θ*_ indicates limit cycles in the network’s phase space that uphold simple periodicity. *γ*-bursts (high *β*_*γ*_) mark heightened attention and learning periods [6, 83]. In our observations, they appear during the mouse’s approach to the reward locations and disappear as it ventures away from them (Fig. 7). Clustering SWR events reflect dense replay activity [55, 56], indicative of periods of memory encoding, retrieval, and network restructurings [57, 84]. Overall, the proposed approach allows studying brain rhythms from a new perspective that complements existing methodology, which may lead to a deeper understanding of the synchronized neuronal dynamics and its physiological functions at temporal mesoscale.

## Acknowledgments

We are grateful to Dr. A. Babichev for fruitful discussions. C.H. and Y.D. are supported by NIH grant R01NS110806 and NSF grant 1901338. C.J. and D.J. are supported by NIH grants R01MH112523 and R01NS097764.

## V. MATHEMATICAL SUPPLEMENT

### Computational algorithms

1. *Kolmogorov score*. Let *X* = *x*_1_ ≤ *x*_2_ ≤ … ≤ *x*_*n*_, be an ordered sequence and let *N*(*X, L*) be the number of elements smaller than *L*,

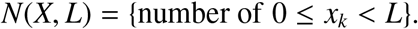

Let 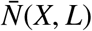 be the expected number of elements that interval (Fig. 9). The closer *X* follows the prescribed behavior, the smaller the normalized deviation^6^

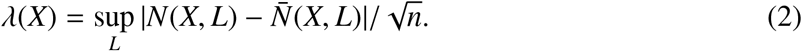

A remarkable observation made in [24] is that the cumulative probability of having *λ*(*X*) smaller than a given *λ* converges to the function

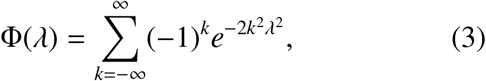

that starts at Φ(0) = 0 and grows to Φ(∞) = 1. The derivative of the cumulative density (3) defines the probability distribution for *λ, P*(*λ*) = ∂_*λ*_Φ(*λ*) (Fig. 2B). Even though the range of *P*(*λ*) includes arbitrarily small or large *λ*s, the shape of the distribution implies that excessively high or low *λ*-values are rare, e.g., sequences with *λ*(*X*) ≤ 0.4 or *λ*(*X*) ≥ 1.8 appear with probability less than 0.3%, Φ(0.4) ≈ 0.003 and Φ(1.8) ≈ 0.997. Since these statistics are universal, i.e., apply to any sequence *X*, the *λ*-score can serve as a universal measure of “stochastic typicality” of a pattern [25–30].
2. *Corrections to Kolmogorov score* up to the order *n*^−3/2^,

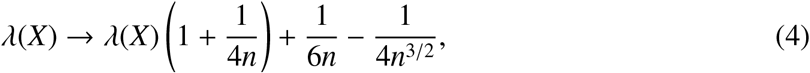

allows for an increase in the accuracy of the finite-sample estimates to over 0.01% for sequences containing as little as 10-20 elements [44–46]. In this study, all *λ*-evaluations are based on the expression (4) and use data sequences that contain more than 25 elements.
3. *Mean Kolmogorov stochasticity score*. The mean *λ* can then be computed as

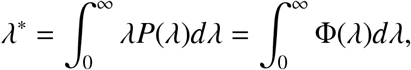

where we used integration by parts and the fact that the distribution *P*(*λ*) starts at 0, *P*(0) = 0, and approaches 0 at infinity, *P*(∞) = 0 (Fig. 2B). Integrating the Gaussian terms in expansion (3) yields Mercator series

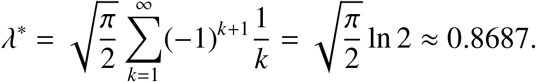
4. Φ(*λ*) *estimates*. For small *λ*s, the Kolmogorov’s Φ-function (3) can be approximated by

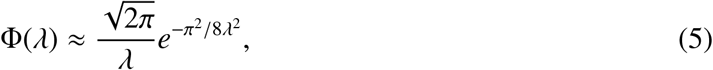

and for large *λ*s, it is approximated by the two lowest-order terms in (3), 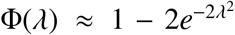 [24, 25]. These formulae allow quick evaluations of the *λ*-scores’ cumulative probabilities outside of the “stochastic typicality band,” *λ <* 0.4 or *λ >* 1.8.
5. *Arnold score*. Let us arrange the points of the sequence *X* on a circle of length *L* and consider the arcs between pairs of consecutive elements, *x*_*i*_ and *x*_*i*+1_ (Fig. 2C). If the lengths of these arcs are *l*_1_, *l*_2_, …, *l*_*n*_, then the sum

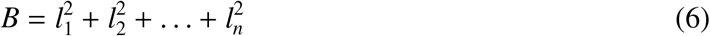

grows monotonically from its smallest value *B*_min_ = *n*(*L/n*)^2^ = *L*^2^*/n*, produced when the points *x*_*k*_ lay equidistantly from each other, to its largest value, *B*_max_ = *L*^2^, attained when all elements share the same location, with the mean *B*^*^ = *B*_min_2*n/*(*n* + 1) 2*B*_min_ [37–40]. Intuitively, orderly arrangements appear if the elements “repel” each other, “clumping” is a sign of attraction, while independent elements are placed randomly. Hence the ratio *β* = *B/B*_min_ can be used to capture the orderliness of patterns:

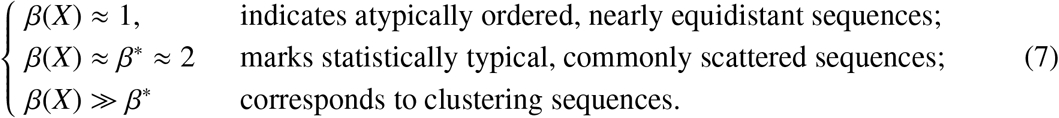
6. *The length L of the circle* accommodating a random sample sequence *X* in Arnold’s method was selected so that the distance between the end points, *x*_0_ and *x*_*n*_, became equal to the mean arc length between the remaining pairs of neighboring points,

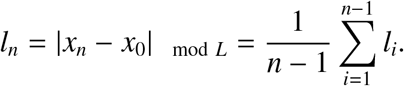
7. *Mean Arnold stochasticity score*. A short derivation of *β*^*^ is provided below for completeness, following the exposition given in [40].
  - The *n* arcs lengths *l*_1_, *l*_2_, …, *l*_*n*_ produced by *n* points, *X* = {*x*_1_, *x*_2_, …, *x*_*n*_}, can be viewed as the “coordinates” of *X* in a *n*-dimensional sequence space. If these coordinates could vary independently on a circle of length *L*, then the sequences would be in one-to-one correspondence with the points of a *n*-dimensional hypercube with the side *L*. However, since the sum of *l*_*i*_s must remain fixed,

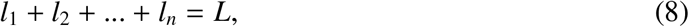

the admissible *l*-values occupy a hyperplane that cuts between the vertices (0, 0, …, 0) and (*L, L*, …, *L*). For example, the two-element sequences described by the coordinates *l*_1_ and *l*_2_ = *L* − *l*_1_ correspond to the points on the diagonal of a *L*-square (Fig. 10A) and three-element sequences correspond to the points of a “diagonal” equilateral triangle in the *L*-cube (Fig. 10B). The four-element sequences are represented by the points of a regular tetrahedron (Fig. 10C) and so forth. Thus, a generic *n*-sequence is represented by a point in a polytope spanned by *n* vertices in (*n* − 1)-dimensional Euclidean space—a (*n* − 1)-*simplex, σ*^(*n*−1)^ [87].
  - The *defining property* of a simplex is that any sub-collection of its vertices spans a sub-simplex: a tetrahedron, *σ*^(3)^, is spanned by four vertices, any three of which span a triangle *σ*^(2)^—a “face” of *σ*^(3)^; any two vertices span an edge, *σ*^(1)^, between them, etc. [87]. Correspondingly, a generic section of the *σ*^(*n*−1)^-simplex by a hyperplane is also a *σ*^(*n*−2)^-simplex (Fig. 10C).
  - Averaging the sum (6) requires evaluating the mean of each 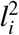,

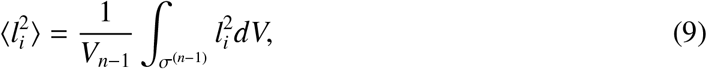

for *i* = 1, 2, …, *n*. Here *V*_*n*−1_ refers to the volume of *σ*^(*n*−1)^ and “*dV*” refers to the volume of a thin layer positioned at a distance *l*_*i*_ away from the *i*^th^ face (Fig. 10C). By the defining property of simplexes mentioned above, the base of this layer is a (*n* − 2)-simplex specified by the equation

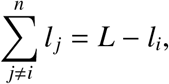

which implies that the sides of this base have length *L* − *l*_*i*_ (just as the sides of *σ*^(*n*−1)^ defined by (8) have length *L*). The volume of the thin layer is *dV* = *C*(*L* − *l*_*i*_)^*n*−2^*dl*_*i*_, so that

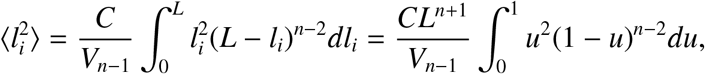

where *u* = *l*_*i*_*/L*. Using the variable *v* = 1 − *u* the latter integral yields:

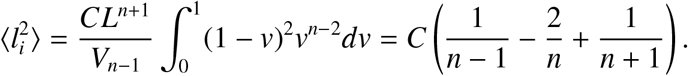

The volume of the *σ*^(*n*−1)^-simplex is

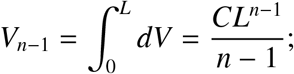

hence the sum (6) divided by *B*_min_ = *L*^2^*/n* yields

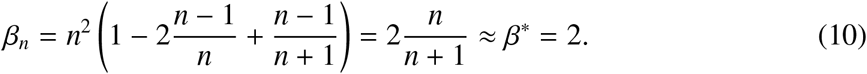
8. *Probability distributions of β-values* form a family parameterized by the number of elements in the sequence. As shown on Fig. 11, these distributions have a well-defined peak at 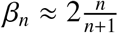 (see below) and rapidly decay as *β* approaches 1 or for *β >* 3.5, which illustrates that typical *β*-values, for all *n*, remain near the impartial mean *β*^*^ ≈ 2.
9. *The sliding window algorithm* can be implemented in two ways:
  - using a *fixed window* that may capture different numbers of events at each step, i.e., *L*_*t*_ = *L*, but *n*_*t*_ and *n*_*t*+1_ may differ;
  - using a *fixed number of events* per window, i.e., *n*_*t*_ = *n*, but *L*_*t*_ and *L*_*t*+1_ may differ.
  In both cases, the mean window width 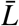 is proportional to the mean separation between nearest events,

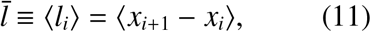

and the mean number 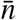 of the data points in the sample sequence, 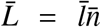. The resulting estimates for *λ*(*t*) and *β*(*t*) are nearly identical and, for qualitative assessments, can be used interchangeably. *10. Local averaging*. To build the dependencies between local averages, we ordered the values assumed by the independent variable, e.g., the speeds, from smallest to largest,

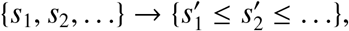

subdivided the resulting sequence into consecutive groups containing 100 elements and averaged each set,

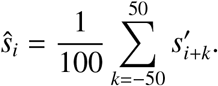

Since each 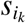 is associated with a particular moment of time 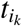, we computed the averages of the corresponding dependent variable, e.g., 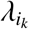,

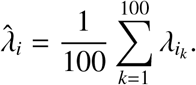

Similarly, ordering the *β*-scores and evaluating their local means produces the 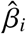-values, along with the means of their 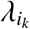-counterparts that occur at the corresponding moments 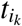, yielding 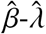 dependence.

**FIG. 9.**
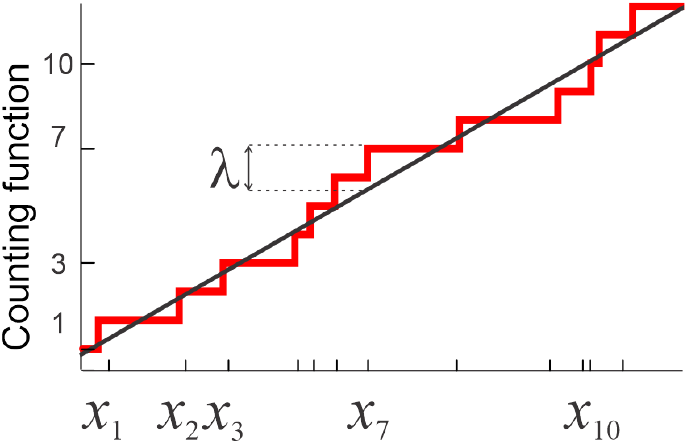
Counting function. *N*(*X*) (red stair-case) makes unit steps at each point of a sequence *X* = {*x*_1_, *x*_2_, …, *x*_*n*_} (tick marks on the *x*-axis). The normalized maximal deviations *λ*(*X*) from the expected mean 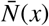 (straight line) exhibit statistical universality and can hence be used for characterizing stochasticity of the individual data sequences *X*.

**FIG. 10.**
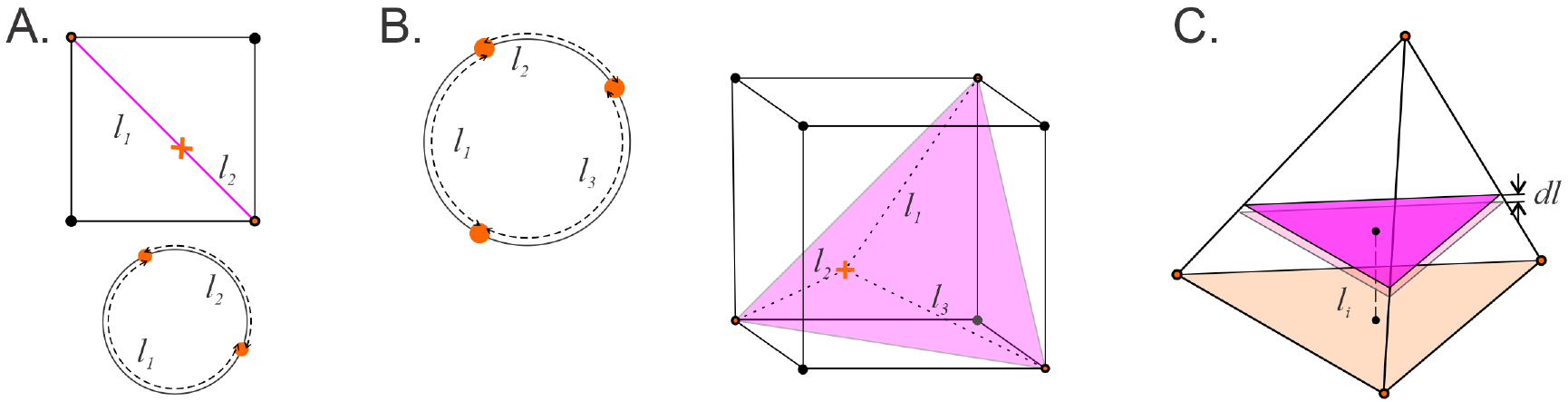
Averaging over a simplex. **A**. If two coordinates *l*_1_ and *l*_2_ of a two-element sequence could independently vary between 0 and *L*, then the pair (*l*_1_, *l*_2_) would cover a 2*D* square. However, if the elements (*x*_1_, *x*_2_) remain on a circle (orange dots below) then the equation (8) restricts (*l*_1_, *l*_2_)-values to the cube’s diagonal (orange cross on the top panel), i.e., to a 1-dimensional simplex. **B**. A configuration of three points on a circle corresponds to a point on the diagonal section of a *L*-cube. **C**. Tetrahedron—a section of a 4*D* cube—is the highest dimensional (3*D*) depictable simplex *σ*^(3)^, which is used to schematically represent *n*-dimensional simplexes, *σ*^(*n*)^. Averaging over 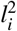 in (9) involves integrating over it the (*n* − 1)-dimensional layers of *σ*^(*n*)^.

**FIG. 11.**
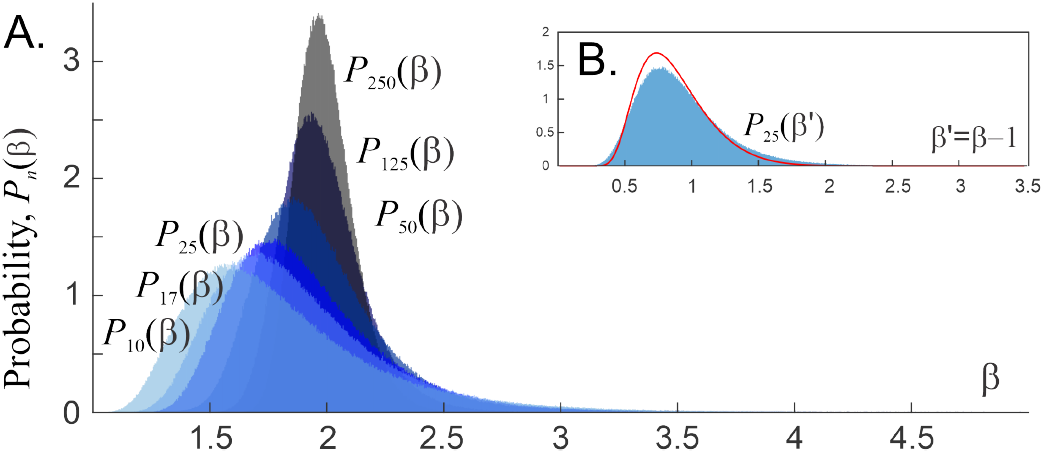
*β*-distributions. **A**. Histograms of *β*-values obtained for 10^6^ sequences containing *n* = 17, 25, 50, 125 and 250 elements peak in a vicinity of the impartial mean *β*^*^ and rapidly decay for *β* ≲ 1.5 and *β* ≳ 3.5. **B**. The distribution of *β*, = *β* − 1 for sequences containing about *n* = 25 points is close to the universal Kolmogorov distribution *P*(*λ*) (red line, Fig. 2A).

## VI. SUPPLEMENTARY FIGURES

**Suppl. Fig. 1.**
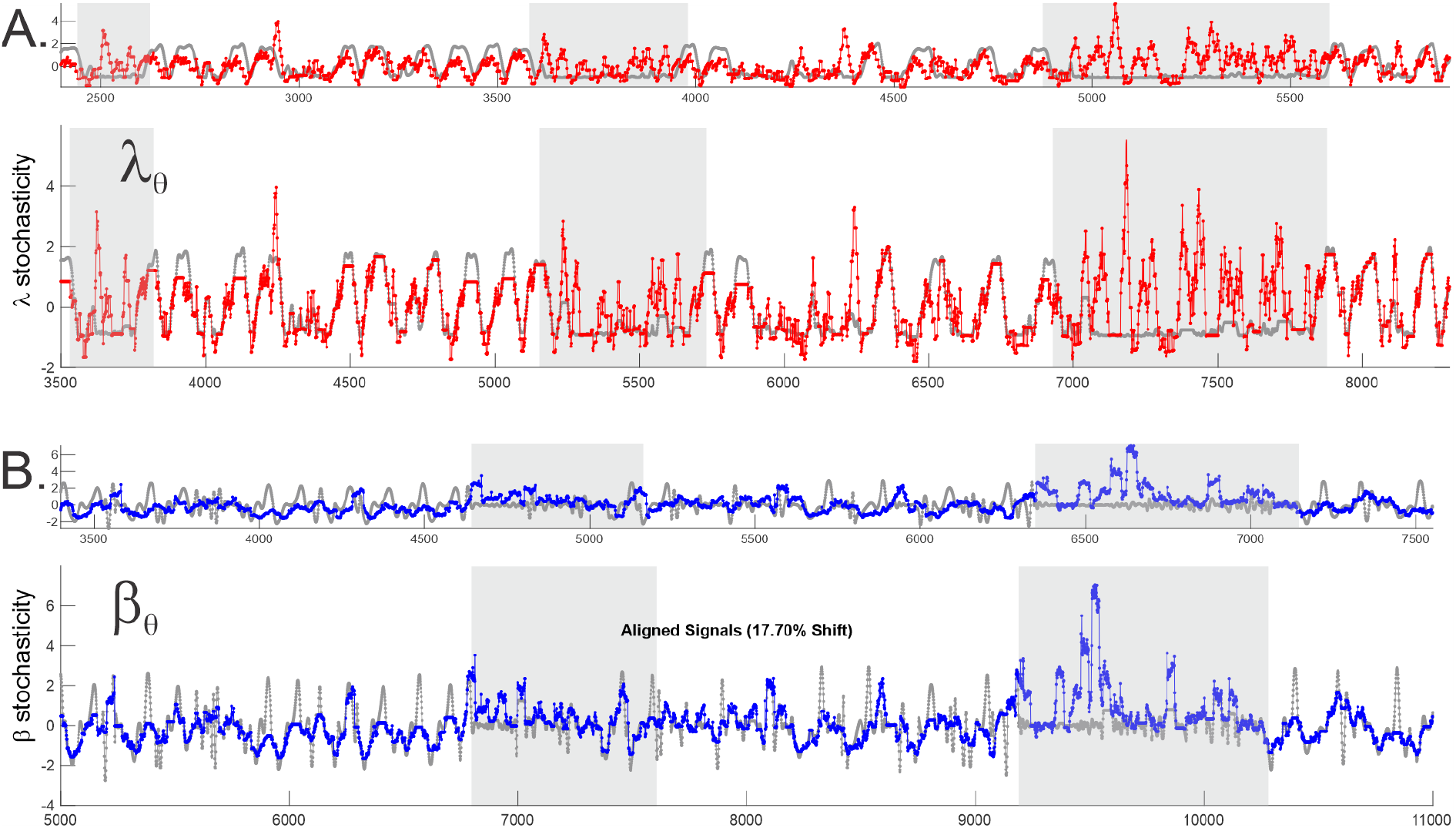
Matching waves using DTW. **A**. Top panels show the original shapes of the speed (*s*(*t*), gray trace) and the *theta*-wave’s Kolmogorov stochasticity score (*λ*_*θ*_(*t*), red trace). Bottom panel shows same functions, matched up by a sequences of local DTW-stretches. Clearly, the speed and the *λ*_*θ*_-stochasticity have the same qualitative shape during active behavior, while during inactive moves (domains marked by light gray stripe) the connection is lost. **B**. Same analyses carried for the mouse’s acceleration (*a*(*t*), gray trace) and Arnold stochasticity parameter (*β*_*theta*_, blue trace). The net amount of stretch required to match speed and *λ*_*θ*_ in this case (including the inactivity periods) is 26%, while the net stretch matching *β*_*θ*_ and the acceleration is ∼ 18%.

**Suppl. Fig. 2.**
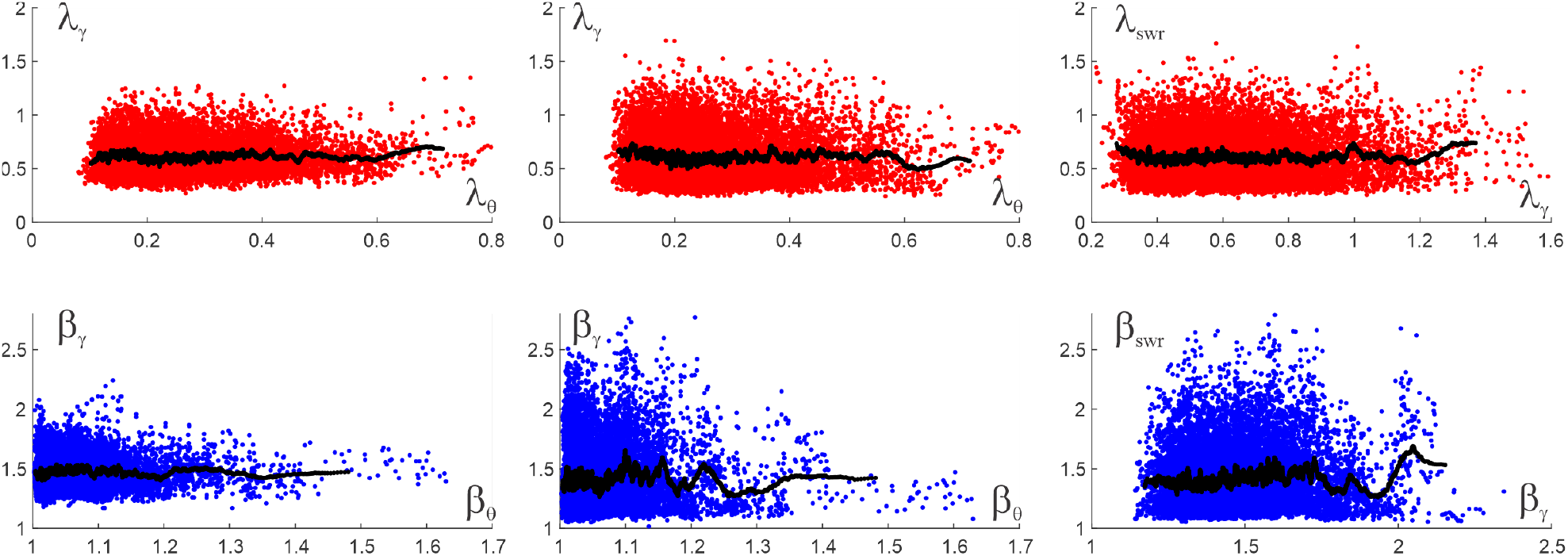
Coupling between stochasticity parameters of different waves. **A**. The geometric layout of points with coordinates (*λ*_*θ*_, *λ*_*γ*_), (*λ*_*θ*_, *λ*_swr_) and (*λ*_*γ*_, *λ*_swr_) shows that a given *λ*-value produced by one wave may pair with any *λ*-value that another wave is capable of producing, i.e., wave patterns deviate from their respective means largely independently from each other. Correspondingly, the locally averaged scores 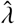 lay approximately horizontally, at the level of the corresponding means ⟨*λ*_*θ*_⟩, ⟨*λ*_*γ*_⟩ and ⟨*λ*_swr_⟩ (see Fig. 4C, Fig. 5B and Fig. 6B). **B**. The *β*-scores reveal similar lack of coupling between waves. Changes in locally averaged 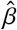-values of one wave not entrain consistent 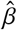-changes of another wave. Thus, the (dis)orderliness of one wave does not enforce the (dis)orderliness of the other and the stochasticity dynamics discussed above provide independent characterizations of of the LFP waves.

## Footnotes

1 Throughout the text, terminological definitions and highlights are given in *italics*.

2 The data used in this work was outlined in [85].

3 DTW separation typically satisfies the triangle inequality, *D*(*a, b*) + *D*(*b, c*) ≥ *D*(*a, c*), which permits interpreting it geometrically, as a distance between signals [49].

4 A succinct expression of this view is provided in [67]: *“rhythmicity is the extent to which future phases can be predicted from the present one*.*”*

5 In his 1955 discussion of the foundations of Quantum Mechanics, John von Neumann attributes a great significance to the fact that *“…the general opinion in theoretical physics had accepted the idea that …continuity …is merely simulated by an averaging process in a world which in truth discontinuous by its very nature. This simulation is such that man generally perceives the sum of many billions of elementary processes simultaneously, so that the leveling law of large numbers completely obscures the real nature of the individual processes*.*”* [68]

6 The supremum, rather than maximum, is required in formula (2) due to discontinuity of the counting function *N*(*X, L*) at the stepping points.

